# TRPV channel OCR-2 is distributed along *C. elegans* chemosensory cilia by diffusion in a local interplay with intraflagellar transport

**DOI:** 10.1101/2020.11.19.390005

**Authors:** Jaap van Krugten, Noémie Danné, Erwin J.G. Peterman

## Abstract

Sensing and reacting to the environment is essential for survival and procreation of most organisms. *Caenorhabditis elegans* senses soluble chemicals with transmembrane proteins (TPs) in the cilia of its chemosensory neurons. Development, maintenance and function of these cilia relies on intraflagellar transport (IFT), in which motor proteins transport cargo, including sensory TPs, back and forth along the ciliary axoneme. Here we use live fluorescence imaging to show that IFT machinery and the sensory TP OCR-2 reversibly redistribute along the cilium after exposure to repellant chemicals. To elucidate the underlying mechanisms, we performed single-molecule tracking experiments and found that OCR-2 distribution depends on an intricate interplay between IFT-driven transport, normal diffusion and subdiffusion that depends on the specific location in the cilium. These insights in the role of IFT on the dynamics of cellular signal transduction contribute to a deeper understanding of the regulation of sensory TPs and chemosensing.

## Introduction

Across the tree of life, organisms depend on avoidance behavior to translocate themselves to a more favorable environment (Behringer et al., 2018; Benov and Fridovich, 1996; Gruntman et al., 2017; Hendry et al., 2018; Moore et al., 2019). In most cases, the ability to sense the outside environment is dependent on transmembrane proteins (TPs) triggering and amplifying downstream signaling pathways, including neuronal activity, that evoke a behavioral response. Primary cilia are slender protrusions of eukaryotic cells that function as signaling hubs, harboring a myriad of TPs involved in signal transduction (Brear et al., 2014; Dwyer et al., 1998; Edwards et al., 2000; Goetz and Anderson, 2010). In general, external stimuli are sensed by G-protein coupled receptors (GPCRs), which relay signals via a series of second messenger molecules like cAMP and calcium to elicit a behavioral or genomic response (Nachury and Mick, 2019; Schou et al., 2015). The ciliary distribution of these TPs is essential for their function, and their location is regulated by the intraflagellar transport (IFT) machinery (Blacque et al., 2004; Mukhopadhyay et al., 2010). Previous work on the dynamics of GPCR’s in mammalian cell lines has shown that they mostly move by passive diffusion, although active transport is observed, most notably during signal-dependent retrieval of GPCR’s out of the cilia (Ye et al., 2018).

The nematode *C. elegans* has proven to be an invaluable model organism to study proteins involved in signal transduction and has been widely used as a model for avoidance behavior and the dynamics of ciliary TPs (Perkins et al. 1986; Culotti and Russel, 1978; Jansen et al. 1999; Hilliard et al. 2004). Particular emphasis has been on *C. elegans* sensory cilia, which are located at the end of dendrites of chemosensory neurons in the amphid and phasmid channels, situated in the head and tail regions of the worm, respectively. Mutagenesis studies, combined with experiments where worms are exposed to a droplet of a chemical compound, have demonstrated that *C. elegans* shows chemotaxis and avoidance behavior and have allowed identification of the neurons involved (Culotti and Russell, 1978; Tran et al., 2017; Ward, 1973). Attractants are sensed by the amphid neurons, while the response to repellents appears to be modulated by a combination of the phasmid neurons and amphid neurons (Hilliard et al., 2002). Together, the sensory neurons and cilia of *C. elegans* allow the nematode to react to environmental stimuli within seconds (Ward, 1973).

Development, maintenance and function of cilia critically depend upon a specific transport mechanism, IFT. IFT consists of transport units called IFT trains, that travel from ciliary base to tip and back again, facilitating transport of ciliary building blocks and sensory proteins (Scholey, 2003). In *C. elegans* cilia, anterograde transport (from base to tip) is driven by heterotrimeric kinesin-II in the first few micrometers of the cilium, after which it gradually hands over the IFT trains to homodimeric kinesin-2 OSM-3 (Prevo et al., 2015; Signor et al., 1999). At the tip of the cilium, IFT trains disassemble and remodel (Mijalkovic et al., 2018). Retrograde transport, back to the base is driven solely by IFT dynein (cytoplasmic dynein-2) (Engel et al., 2012; Mijalkovic et al., 2017; Pazour et al., 1999).

The backbone of these trains is formed by the IFT-A, IFT-B and BBSome particle complexes. The link between IFT trains and functional cargo like GPCRs is most likely formed by Tubby proteins and the BBSome. Tubby-like protein 3 (TULP3), together with IFT-A, was found to be essential for the ciliary import of some GPCRs (Badgandi et al., 2017; Mukhopadhyay et al., 2010). Mutations in the BBSome and the *C. elegans* TULP3 homolog, tub-1, show chemotaxis defective behavior, demonstrating the importance of the localization and active transport of sensory TPs (Blacque et al. 2004, Mukhopadhyay et al., 2005).

Electron microscopy and experiments with fluorescent beads tethered to membrane glycoproteins in *Chlamydomonas* have shown directly that IFT trains are physically linked to the membrane (Pigino et al., 2009; Shih et al., 2013). Studies in mouse, *Chlamydomonas* and *C. elegans* have demonstrated that, among other cargoes, several types of TPs are transported by IFT trains (Lechtreck, 2015). Ensemble fluorescence microscopy studies have shown movement by IFT of the ion-channels PKD2 and OCR-2, in *Chlamydomonas* and *C. elegans* respectively (Huang et al., 2007; Qin et al., 2005). Elegant single-molecule studies in cultured mammalian cells demonstrate ciliary TPs exhibiting complex dynamics of both passive diffusion and active transport by IFT (Milenkovic et al., 2015; Weiss et al., 2019; Ye et al., 2013). While occasionally undergoing active transport, ciliary receptors like Smoothened were found to be frequently confined at the ciliary base, and to mostly exhibit passive diffusion in the ciliary membrane.

Despite this progress on the understanding of the transport mechanisms that transport TPs in cilia many questions remain. It is, for example, unclear what the interplay is between active, IFT-driven, and passive, diffusive transport of TPs, how this is regulated and whether it depends on stimulation of the cilia. To shed light on this, we investigate here the dynamics underlying the redistribution of ciliary components during avoidance behavior in the chemosensory cilia of live *C. elegans* using sensitive fluorescence microscopy. With the goal to connect the whole-organism avoidance behavior to the single-molecule dynamics of the proteins at the basis of sensory signal transduction. We find that repellent addition results in a reversible redistribution of the IFT machinery, the ciliary axoneme, and the TRPV cation channel OCR-2 out of the ciliary distal segment (DS). To elucidate the dynamics underlying this redistribution, we performed single-molecule fluorescence microscopy of OCR-2 in the phasmid cilia of live *C. elegans*. Analysis of the trajectories reveals that active, IFT-driven transport is the predominant mode of transport in the dendrite and transition zone. Along the cilia, the proper distribution of OCR-2 depends on an intricate location-specific interplay between active transport, normal diffusion and subdiffusion, with the latter playing an important role in the confinement of TPs at the ciliary tip. These insights in the dynamics of cellular signal transduction during external stimulation contribute to a wider understanding of IFT dynamics and to cilia as chemosensory organelles triggering behavioral responses.

## Results and discussions

### Exposure to repellents leads to reversible redistribution of IFT components

Exposing *C. elegans* to certain chemicals, including SDS, leads to a drastic change in behavior: the worm abruptly ceases its elegant, sinusoidal motion and changes direction. This behavior is called the avoidance response and is triggered by its sensory cilia (Perkins et al., 1986). The nervous system of *C. elegans* consists of 302 neurons, of which 32 are ciliated. Most of these cilia are situated in the two amphid openings, in the head region of the nematode. Because of their number, and the high background fluorescence around the amphid cilia, we study the two pairs of phasmid cilia (PHAL, PHAR, PHBL and PHBR), located in the tail of the animal. To obtain insight in the molecular response of the phasmid chemosensory cilia to aversive compounds, we added water-soluble repellant chemicals while imaging the cilia using epi-illuminated widefìeld fluorescence microscopy. Strains endogenously expressing fluorescently labeled TBB-4 (TBB-4::EGFP) as marker for axonemal microtubules and XBX-1 (XBX-l::EGFP) (Mijalkovic et al., 2017), an IFT-Dynein marker that allows probing IFT dynamics, were exposed to 0.1% (W/V) sodium dodecyl sulfate (SDS), by pipetting a droplet on them. SDS is thought to be a chemical mimic of dodecanoic acid, a compound secreted by nematicide-secreting *Streptomyces* bacteria (Tran et al., 2017). *C. elegans* has been demonstrated to avoid SDS, even at a concentration as low as 0.1%, which is below its critical micelle concentration (Rahman and Brown, 1983). To test the spreading of SDS, the fluorescent compound fluorescein was added in an identical way. This experiment indicates that nematodes, when exposed to a droplet of dissolved chemical compound, experience an immediate increase of compound concentration, which settles to a maximum concentration after two minutes (Figure S1).

Even though worms had been anesthetized with 5mM Levamisole, they still exhibited an avoidance response within seconds after SDS addition, as evident from a sudden movement of the organisms (Movie SI). We set this particular moment in time to zero. This movement, together with the adverse effect of droplet addition on imaging, made constant refocusing necessary, rendering the frames of the first tens of seconds after addition impossible to interpret in most cases. To visualize the distribution of the fluorescent ciliary components, we time-averaged (over 3 s) subsequent fluorescence images such as the one shown in (Figure 1A, Movie S1). First, we focus on IFT Dynein. The images show that, before SDS addition, IFT dynein occupies the entire length of the phasmid cilia, as observed before (Mijalkovic et al., 2017). After about a minute, the motors have drastically redistributed, extending only from the base of the cilium up to the end of the middle segment. At longer time scales the distribution appears to have recovered completely to the original situation, before applying the stimulus.

**Figure 1:**
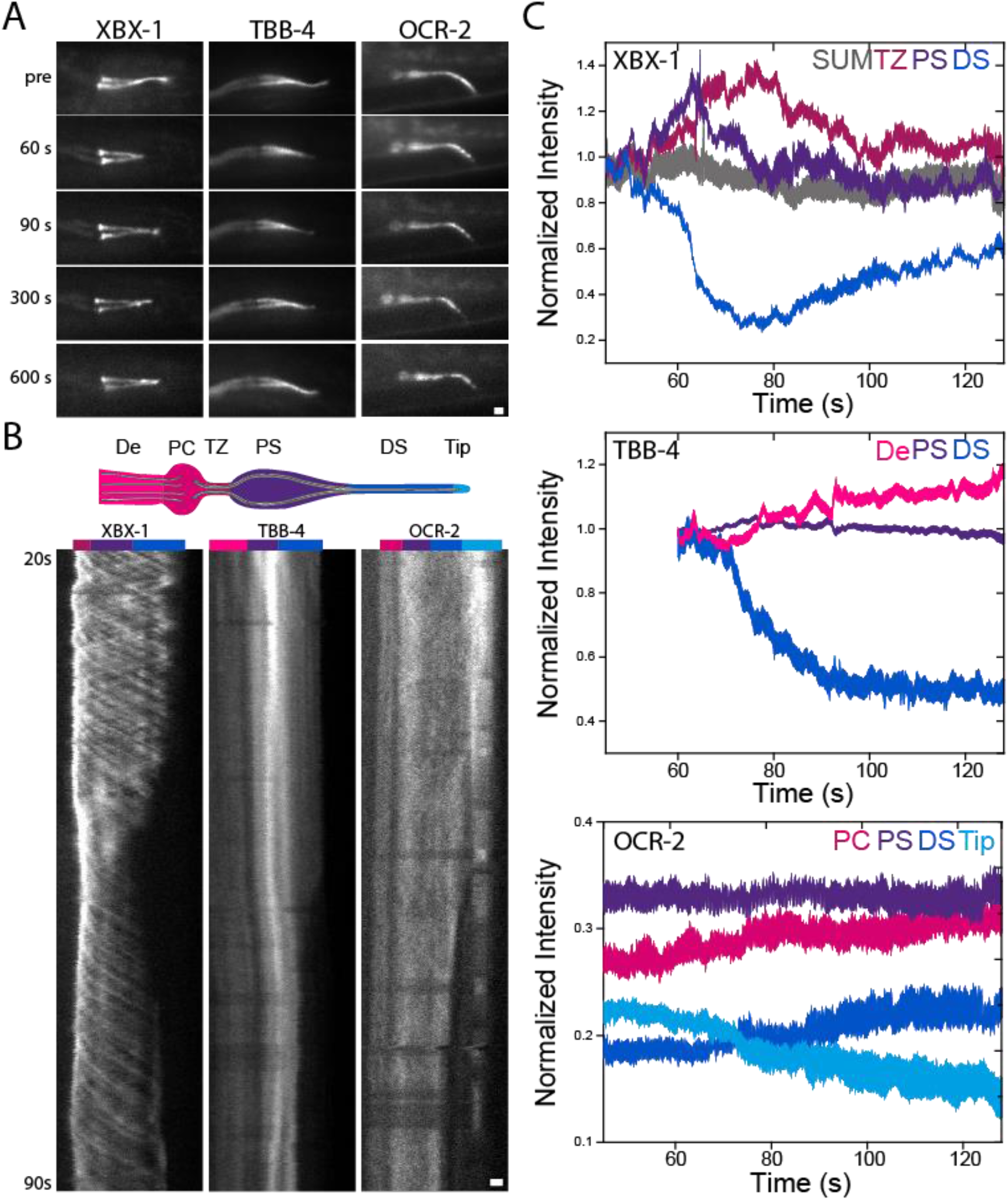
Ciliary components redistribute reversibly upon SDS exposure. **(A)** Time-averaged (over 3 s) fluorescence images of TBB-4::EGFP, XBX-l::EGFP and OCR-2::EGFP before and after SDS exposure. Scale bar: 1 μm. **(B)** Cartoon illustrating the ciliary segments (PCMC: periciliary membrane compartment; TZ: transition zone; PS: proximal segment; DS: distal segment) and representative kymographs of TBB-4::EGFP, XBX-l::EGFP and OCR-2::EGFP starting 20 s after SDS exposure. Scale bar: 1 μm. (C) Relative fluorescence intensities of TBB-4::EGFP, XBX-l::EGFP and OCR-2-EGFP in different ciliary segments. For TBB-4::EGFP and XBX-l::EGFP, normalized by the average fluorescence intensities of the first 10 frames. For OCR-2::EGFP, the total fluorescence intensity along the whole cilium was normalized, immediately after SDS exposure.

To get more insight in the redistribution dynamics, we generated kymographs from the original fluorescence image sequences (Mangeol et al., 2016) (Figure 1B). The colored bars at the top of the kymographs indicate the dendrite, PCMC and ciliary segments, as schematically represented in the cartoon on top. The kymograph shows that, up to about a minute after SDS exposure, processive IFT-dynein trains can be observed moving from ciliary base to tip. After a minute, however, IFT dynein suddenly redistributes out of the distal segment (DS), no longer reaching the ciliary tip, but recovers its pre-exposure distribution quickly after.

To quantitatively compare the amounts of proteins in the different ciliary segments in several worms, the average fluorescence intensity per segment is shown as a function of time (Figure 1C; color coding of the ciliary segments as in cartoon). These time traces show that the amount of IFT dynein in the DS rapidly (within −20 seconds) decreases about five-fold, −1 minute after SDS addition. The traces indicate that IFT dynein moves away from the distal segment (DS), through the proximal segment (PS), towards the transition zone (TZ) and base (Figure 1B and C). The total amount of IFT dynein does not seem to change, indicating that the motors do not leave the cilium on this time scale. Whereas the avoidance reaction of the worms occurs within seconds after SDS addition, IFT dynein reacts later, after 49.7 ± 4.7 s (mean ± s.e.m.; n = 8 worms). This remarkable delay, between avoidance response and IFT-dynein redistribution, will be discussed further below. Taken together these result show that, about a minute after exposure to SDS, IFT dynein reversibly redistributes, temporarily leaving the DS towards the PS and ciliary base.

IFT is required for the maintenance of the cilium, and transports, among others, tubulin towards the tip of the ciliary axoneme (Kozminski et al., 1993; Marshall and Rosenbaum, 2001). To test whether the depletion of IFT dynein in the distal segment after SDS exposure leads to axoneme instability, we next imaged worms expressing fluorescent tubulin. The time-averaged fluorescence images and kymograph of TBB4::GFP (Figure 1A, B and C) show a similar redistribution as observed for IFT dynein, albeit slightly later, 68.6 ± 6.7s (mean ± s.e.m.; n = 7 worms)). This disappearance of TBB-4 from the end of the cilium is most likely caused by shortening of the ciliary axoneme. Remarkably, the shortening appears to stop at the end of the PS, where the axoneme consists of microtubule doublets. In a previous study, femtosecond-laser ablation of the phasmid-neuron dendrites was shown to also cause the collapse of the DS, but not the PS, suggesting that the PS, consisting of microtubule doublets, is more stable than the DS, which consists of microtubule singlets (Mijalkovic et al., 2020). In contrast to IFT dynein, part of the tubulin appears to leave the cilium, as is evident from the increased TBB-4 fluorescence in the dendrite (Figure 1C). The −19 s delay between the onset of TBB-4 and IFT dynein redistribution might indicate that axoneme shortening is a consequence of the redistribution of the IFT machinery resulting in depletion of free tubulin at the axonemal tip. On longer time scales, the TBB4::GFP fluorescence distribution does not show as clearly a recovery as IFT dynein.

To visualize the distribution of TPs involved in chemosensation and their dynamics during avoidance behavior, we generated worms endogenously expressing the fluorescently labeled GPCR SRB-6 (SRB-6::EGFP) and transmembrane calcium channel OCR-2 (OCR-2::EGFP) (Tobin et al., 2002; Tran et al., 2017). Expression levels of SRB-6::EGFP were low, only several tens of molecules could be observed in each phasmid cilia pair (Figure S2, Movie S2). Such expression levels are too low to perform repellent experiments, which require long-term imaging without photobleaching. High laser powers are required to visualize such a small number of molecules, resulting in bleaching of all present molecules within seconds. Expression levels of OCR-2, on the other hand, were much higher, making it possible to image OCR-2 long enough at laser powers low enough to avoid substantial photobleaching during the course of the experiment.

To probe whether the redistribution of IFT dynein and the partial shortening of the axoneme caused by SDS has an effect on the distribution of TPs like OCR-2, we performed SDS-addition experiments with worms expressing OCR-2::EGFP. Time-averaged fluorescence images show that, before application of SDS, OCR-2 is distributed unevenly over the cilium (Fig 1A): it appears to accumulate at the base, in the DS and the tip, while it is less abundant in the middle of the cilium and almost absent in the TZ (see also below). After SDS addition, OCR-2 redistributes, becoming less abundant in tip and DS, but to a different extend than IFT dynein and tubulin. Kymographs showing the OCR-2::EGFP distribution confirm the observation that OCR-2 leaves the DS (Figure 1B). Part of the OCR-2 remains, indicating that the ciliary membrane does not retract from the DS (together with the microtubules). For further quantification of the changes in fluorescence intensity, we divided the cilia in four segments, as indicated on top of the kymograph (including the ciliary tip as distinct segment). The fluorescence intensity time traces in these four different segments show that, while the amount of OCR-2 in the PS remains more or less constant, approximately one third of the proteins in the tip disappears, and an equal amount reappears in the DS and PCMC (Figure 1C). Together, these data show that the cilium is capable of rapid redistribution and recovery of IFT in the distal segment, resulting in a partial shortening of the axoneme and a reduction in the amount of OCR-2 in the tip region.

Rapid redistribution of IFT dynein and tubulin away from the DS is reminiscent of the laser ablation and azide dosing experiments we have performed before (Mijalkovic et al., 2020). Here it was shown that ablation of the phasmid chemosensory neurons at the dendrite (a couple of micrometers before the start of the cilium), resulted in the activation of retrograde IFT, leading to a redistribution of IFT components out of the DS towards the base on a time scale similar to the SDS-addition experiments presented here. ATP did not seem to be involved in the initial response, since ATP depletion by azide-addition did not lead to the activation of retrograde IFT. Instead, upon depletion of ATP, IFT components slowed down, redistributed towards the base and eventually left the cilium, comparable to the response to laser ablation at longer time scales. Together, these data were interpreted to reveal an IFT-regulatory mechanism that allows rapid redistribution of ciliary components in response to changes in the outside environment. In the present SDS-addition experiments we see qualitatively similar redistributions of tubulin and IFT components, with one crucial difference: the redistribution starts about one minute after the SDS addition and almost concomitant avoidance response (within a few seconds), while redistribution starts within a few seconds after laser ablation or azide-addition. So, although immediate redistribution is possible in the cilium, it seems that upon SDS addition, the trigger for redistribution is delayed. Since the response time for second messengers is known to be within seconds, they are not likely the cause of the delay (Jiang et al., 2019; Pifferi et al., 2006; Tran et al., 2017). In our experiments, the worms are exposed to a prolonged SDS stimulus. This might indicate that the localization of sensory TPs affects cilium sensory sensitivity and that the redistribution effect observed here is a habituation response.

### Ensemble distribution and dynamics of OCR-2

We next had a closer look into the distribution of OCR-2 along the cilium. First, time-averaged fluorescence images were generated from image sequences of a *C. elegans* strain expressing OCR-2-EGFP (Figure 2A-D). In order to localize the TZ with respect to the OCR-2 distribution, we imaged phasmid cilia in worms overexpressing the TZ marker MKS-6::mScarlet and with endogenously labeled OCR-2 (Figure 2C-D). From such images, the intensity distribution along the cilium long axis was calculated (Figure 2E). The images and intensity distribution indicate that the TZ corresponds to a region with very low OCR-2 density. Anterior to the TZ, OCR-2 appears to accumulate in a cup-like shape, which most likely corresponds to the PCMC. Posterior to the TZ, the intensity of OCR-2 is relatively high, dropping further along the middle segment and increasing again in the DS, peaking at the ciliary tip. A super-resolution image obtained from many localizations of individual OCR-2 molecules (see below and Oswald et al., 2018), confirms these observations (Figure 2B), highlighting the membrane localization of OCR-2, in particular in the wide PS.

**Figure 2:**
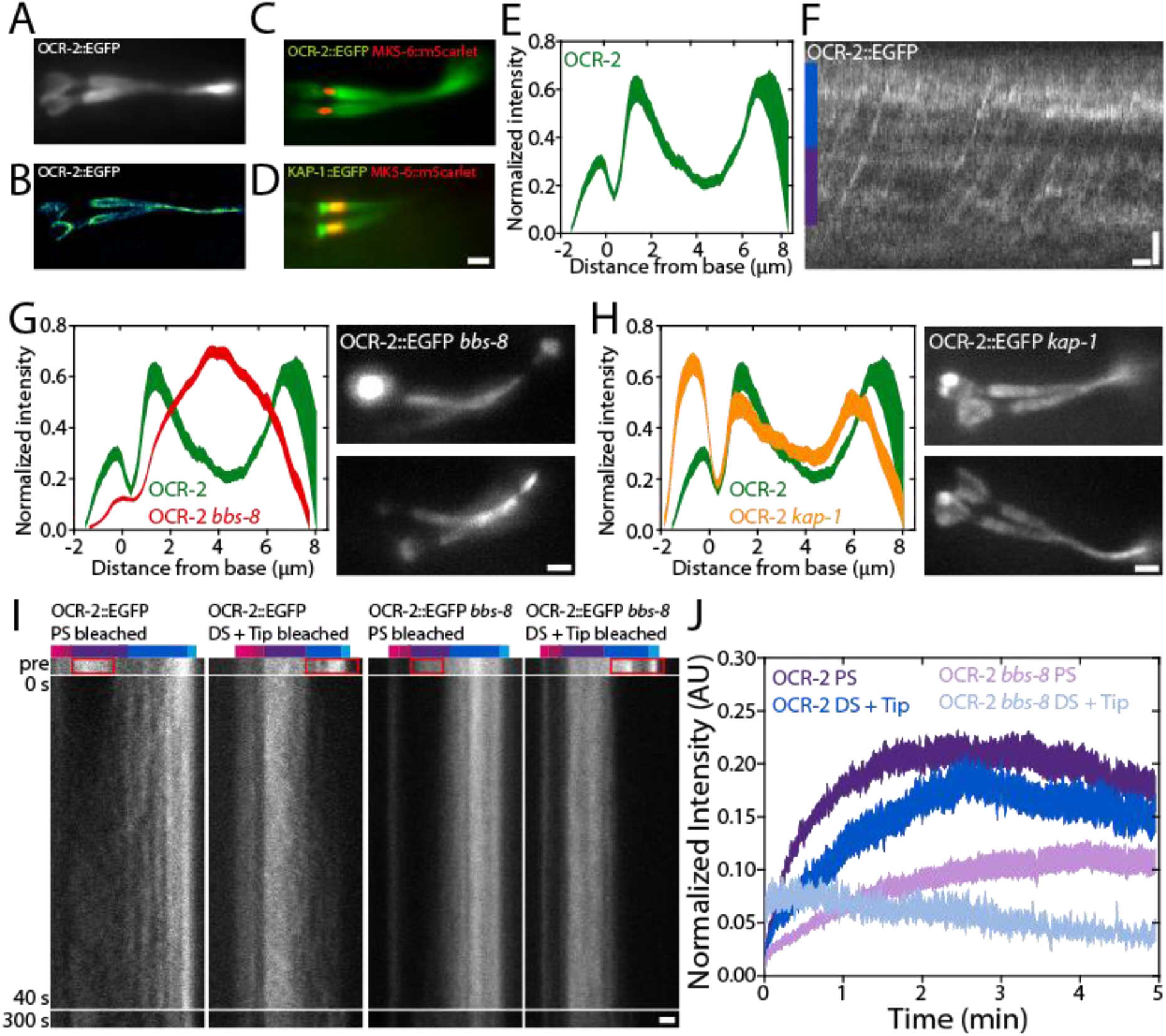
Ensemble distribution and dynamics of OCR-2. **(A)** Time-averaged (over 3 s) fluorescence images of OCR-2::EGFP. **(B)** Super-resolution image of OCR-2::EGFP molecule localizations. **(C)** Dualcolor image of endogenously labeled OCR-2::EGFP and overexpressed MKS-6::mScarlet. (D) Dual-color image of endogenously labeled Kinesin-II(KAP-l::EGFP) and MKS-6::mCherry. **(E)** Average intensity distribution of OCR-2::EGFP along the cilium (n = 22 worms, normalized intensity distributions). **(F)** Kymograph of ensemble dynamics of OCR-2::EGFP with bars indicating the PS (purple) and DS (dark blue). Scale bars: 1 μm and 1 s. (G) Left: normalized average intensity distribution of OCR-2::EGFP (green) and OCR-2::EGFP *bbs-8* (red) (n = 71) along the cilium. Right: Time-averaged fluorescence images of OCR-2::EGFP *bbs-8*. **(H)** Left: normalized average ciliary distribution of OCR-2::EGFP (green; same data as E) and OCR-2::EGFP, kap-1 mutant (red) (n = 26 worms). Right: Time-averaged (over 3 s) fluorescence images of OCR-2::EGFP, kap-1 mutant. Scale bar: 1 μm. (I) Representative kymographs before and after photo bleaching the PS, or DS and Tip of OCR-2::EGFP and OCR-2::EGFP *bbs-8*. Colored bars indicate segments as in Figure 1B. Time scale is identical for all kymographs. (J) Normalized average intensity of OCR-2::EGFP in PS, or DS and Tip, after photobleaching PS or DS and Tip respectively, in WT and *bbs-8* animals (OCR-2::EGFP PS: n = 11, DS = Tip: n =10; OCR-2::EGFP *bbs-ĩ>* PS: n = 16, DS + Tip: n = 10).

To get a first insight in what underlies the distribution of OCR-2 along the cilium, we generated kymographs from the original image sequences at the ensemble level (Figure 2F). The kymograph clearly shows directional, processive movement of OCR-2, in agreement with previous studies of worms expressing fluorescent OCR-2 from extra-chromosomal arrays (Qin et al. 2005). The continuous lines due to motor-driven OCR-2, however, are less clear than what we have observed before for motor proteins and other IFT components (Mijalkovic et al., 2017; Prevo et al., 2015) and appear to be obscured by a relatively high, unstructured background signal of OCR-2. This might indicate that OCR-2 is only partly actively transported by IFT and to a substantial account by passive diffusion.

In order to determine how essential active transport by IFT is for the ciliary distribution of OCR-2, we generated worms lacking BBS-8, which is crucial for a functional BBSome, and thereby for the connection between ciliary membrane proteins and the IFT machinery (Blacque et al., 2004). Kymographs generated from image sequences, indeed, do not show any signs of active transport of fluorescent OCR-2 in the absence of BBS-8 function, but do show that IFT is still active (Figure S3). Fluorescence images were time averaged in order to generate an intensity distribution averaged over multiple cilia (Figure 2G). These images show that OCR-2 does enter the cilia of *bbs-8* worms, but that its distribution along the cilium is remarkably different compared to wild-type animals: OCR-2 concentration does not peak behind the TZ and at the tip (as found in wild type), but shows a maximum in the middle of the cilium. We note that the time-averaged fluorescence images of *bbs-8* worms were quite heterogeneous. In some worms, substantial accumulations of OCR-2 were observed in the PCMC (e.g. bottom image in Figure 2G) suggesting that import of OCR-2 is severely hampered in these cilia. Together, these results show that active transport plays an important role in the ciliary distribution of the TP OCR-2. Furthermore, although a functional BBSome has been shown to be necessary for avoidance behavior and is involved in ciliary exit of TPs (Blacque et al., 2004; Lechtreck et al., 2013), it seems not completely essential for ciliary entry of OCR-2 into the cilium. Together, this might indicate that the proper, IFT-dependent, ciliary distribution of TPs involved in chemotaxis is required for the propagation of environmental stimuli.

To study the effect of a more subtle modulation of active transport of TPs, we generated worms with endogenously labeled OCR-2, but lacking Kinesin-II function (tml23). Kinesin-II drives, together with OSM-3, anterograde IFT, with Kinesin-II mostly active from the ciliary base across the TZ and OSM-3 in the rest of the cilium (Prevo et al., 2015). While mutant worms lacking OSM-3 function have short cilia lacking a DS and are osmotic avoidance defective, worms lacking Kinesin-II function do show osmotic avoidance behavior and exhibit only minor structural defects (Oswald et al., 2018; Snow et al., 2004). Time-averaged image sequences and the average intensity distribution of OCR-2 in kap-1 mutant cilia show accumulations of OCR-2 posterior to the TZ and at the ciliary tip, just as wild type, but also show a far more pronounced accumulation in the PCMC (Fig 2J). These observations are in line with earlier observations of the distributions of IFT-particle components in kap-1 mutant strains, which showed similar distributions along the cilium compared to wild type, but also showed accumulations in the PCMC, suggesting that Kinesin-II is a more efficient import motor than OSM-3 (Prevo et al., 2015). Furthermore, these observations are in line with those of OCR-2 in bbs-8 mutant worms, supporting the conclusion that efficient, active transport of OCR-2 by IFT is essential for its distribution and function.

To obtain insight in the ensemble dynamics that underlie the ciliary distribution of OCR-2, we performed Fluorescence Recovery After Photobleaching (FRAP) experiments on OCR-2::EGFP in wild-type and bbs-8 mutant worms (Fig 2I), photobleaching the EGFP in PS, or DS and tip, by brief, intense laser illumination. Image sequences of the fluorescence recovery were converted into kymographs of the cilium (Fig 2I) and fluorescence intensity time traces integrated over the ciliary segments (Fig 2J). Kymographs and time traces show that, in wild-type cilia, the rate and extend of OCR-2::EGFP recovery in the PS after bleaching the PS, is slightly different from the recovery in DS and Tip after bleaching DS and Tip. In the PS, only −21% of the intensity before bleaching the PS recovered after −2 minutes, while in the DS and Tip −18% recovered −2 minutes after bleaching DS and Tip. This indicates that in the cilium, only a small fraction of OCR-2 is free to move, driven by IFT or free diffusion.

In bbs-8 mutant worms (where the connection between OCR-2 and IFT machinery is disrupted), the mobile fraction is even smaller in the PS (−11%) and DS (−5%) compared to those segments wild type. Compared to wild type, a similar difference in the recovery rate and extend between the PS and DS in the bbs-8 mutant cilia was observed. The difference of the mobile fractions in the PS and DS between wild type and *bbs-8* mutant worms furthermore demonstrates the importance of IFT for the ciliary distribution of OCR-2. This difference is larger in the DS, suggesting IFT plays a larger role in the DS for OCR-2 localization.

Collectively, these findings demonstrate the diversity of transport modalities that underlie the ensemble dynamics resulting in the steady-state distribution of OCR-2 along the cilium. This distribution is not uniform along the cilium but shows accumulations at the Tip and posterior to the TZ, and a smaller accumulation anterior to the TZ. In mutant animals with perturbed active transport of OCR-2, the distribution is different. In addition, our imaging and FRAP analysis indicate that diffusion of OCR-2 in the ciliary membrane also plays a key role.

### Single-molecule imaging reveals large diversity in OCR-2 motility

In order to obtain more detailed insight into the extents by which diffusion and active transport drive OCR-2 motility along the cilium, we performed single-molecule tracking of OCR-2::EGFP in live *C. elegans* using laser-illuminated wide-field epifluorescence microscopy, at a frame rate of 10 Hz. In Figure 3A (see also Movie S3), example trajectories are represented as kymographs, with location in the cilium indicated. The kymographs suggest a variety of different behaviors of OCR-2, ranging from diffusion to directed transport in anterograde and retrograde directions. The specific behavior appears to depend on the actual location in the cilium. In the dendrite, straight, slanted lines are observed towards the PCMC (see also Figure S4) (1), indicative of active, directional transport of OCR-2. In the PCMC, kymograph lines appear mostly horizontal but slightly wavy (2), indicative of particles lingering a substantial amount of time (which appears to be limited by photobleaching in our experiments). In the TZ, again straight, slanted lines are observed (3), indicating that particles cross the TZ mostly by directed transport. In the PS, more heterogeneous behavior is observed. Some particles show straight, slanted lines (4), indicative of active transport, while others appear saltatory (5), indicating that particles diffuse or move by short bouts of active transport in anterograde or retrograde direction. In the DS, a similar pattern of directed motion in combination with saltatory movement is observed (6), albeit that the fraction of directed transport appears higher. Taken together, single-molecule kymographs hint at a large diversity of OCR-2-motility patterns that appear to depend on the specific ciliary segment and location.

**Figure 3:**
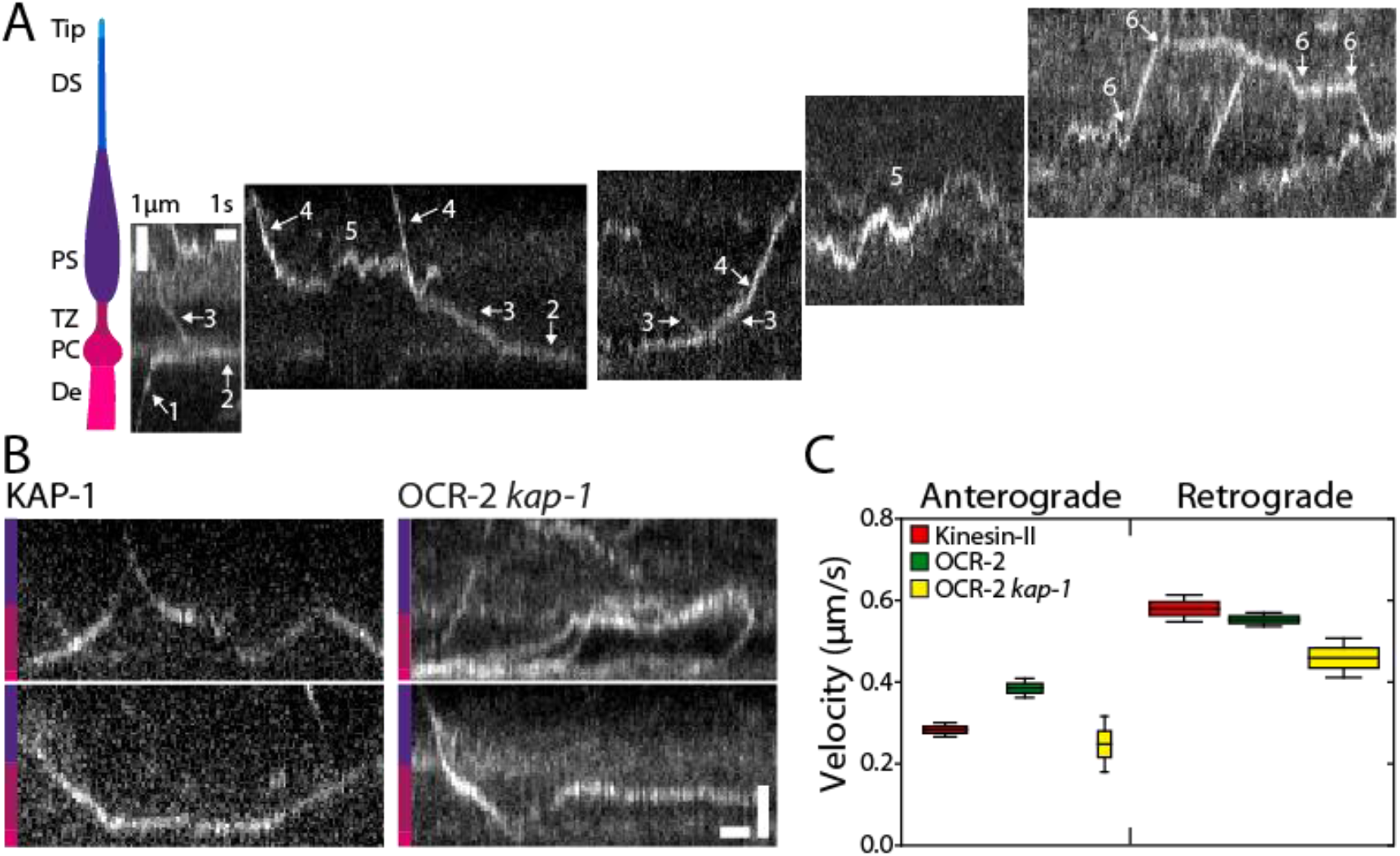
Single-molecule imaging of OCR-2. **(A)** Cartoon of the ciliary segments and representative single-molecule kymographs of OCR-2::EGFP. Scale bars, vertical: 1 μm, horizontal: 1 s. (B) Singlemolecule kymographs showing TZ crossings of Kinesin-II (KAP-l::EGFP) and OCR-2::EGFP *kap-1*(tml23). Color bars indicate ciliary segment as in (A). Scale bars, vertical: 1 μm; horizontal: 1 s. **(C)** Anterograde (A) and retrograde (R) velocity of KAP-l::EGFP (A: 63 trajectories; R: 273 trajectories), OCR-2::EGFP (A: 51 trajectories; R: 297 trajectories) and OCR-2::EGFP *kap-1*(tml23) (A: 19 trajectories; R 141 trajectories) (error bars represent s.e.m.).

One of the observations in the kymographs of Fig 3A is that the motion of OCR-2 in the TZ appears to be mostly directional, driven by IFT. The motility of cargo OCR-2 appears very similar to that of motor Kinesin-II (imaged using a KAP-l::EGFP strain; Figure 3B), also with comparable velocity in anterograde and retrograde direction (Figure 3C). These observations strongly suggest that OCR-2 is indeed transported across the TZ by IFT trains driven by Kinesin-II in the anterograde direction and IFT dynein in the retrograde direction. The densely packed protein network (e.g. the Y-shaped linkers connecting axoneme and membrane) of the TZ forms a diffusion barrier between dendrite and PS (Garcia-Gonzalo and Reiter, 2017). We have shown before that in the absence of Kinesin-II function (in the kap-1 mutant strain), OSM-3 can take over and drive IFT across the TZ. In these strains, however, the frequency and number of IFT trains entering the PS appears affected and IFT components show more substantial accumulations in front of the TZ (Prevo et al., 2015). In line with these observations, we have shown above that OCR-2 is still present in cilia lacking kinesin-II function, but that its ciliary distribution is affected (Figure 2H). Single-molecule kymographs generated from image sequences of kap-1 mutant worms show many more OCR-2 molecules being stuck in the TZ (Figure 3B), or traversing it more irregularly than in wild-type animals, although successful TZ crossings can still be observed. Velocities extracted from the trajectories were only slightly lower than for wild type (Figure 3C), which might seem remarkable, since OSM-3, which is generally considered to be a faster motor, drives anterograde crossing (in the kap-1 mutant) instead of Kinesin-II (in wild type). We have, however, shown before that OSM-3 velocity in the TZ of kap-1 mutant worms is substantially reduced in addition to affecting OSM-3’s efficiency to traverse the TZ, most likely due to TZ structures acting as roadblocks for OSM-3, more severely than for Kinesin-II (Prevo et al., 2015). Taken together, the TZ acts as an efficient diffusion barrier for OCR-2, which needs to be overcome by active transport driven by IFT. In mutant worms lacking Kinesin-II function, OCR-2 transport over the TZ can be taken over by OSM-3, albeit with loss of efficiency. In both wild-type and kap-1 mutant worms, IFT dynein drives effective retrograde transport of OCR-2.

### Single-molecule imaging reveals bimodal motility distribution of ciliary OCR-2

To get a more quantitative insight in the motility behavior of single OCR-2 molecules inside the entire cilium, we tracked single molecules in images sequences, such as those represented in the kymographs of Figure 3A, using a local ciliary coordinate system (parallel and perpendicular to a spline drawn along the long axis of each cilium). Representative trajectories are shown in Figure 4A, with examples of a trajectory suggestive of active, directed transport (*), a trajectory suggestive of diffusive transport (+) and a trajectory of a molecule that hardly moves in the ciliary tip (#). For further analysis, we calculated the mean squared displacement (MSD) of these example trajectories (Figure 4B&C). MSD analysis is a widely used approach to characterize motility, based on the scaling of MSD with time (Gal et al., 2013). This scaling can be expressed in one dimension as: *MSD*(*τ*) = 2·Γ·*τ^α^*, with, Γ the generalized transport coefficient, *τ* the time lag, and *α* the exponent. In case *α* = 2, the motion is purely directed (ballistic) and the velocity (*v*) is equal to 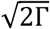. When *α* = 1, the motion is due to normal diffusion and characterized by a diffusion coefficient (*D*) equal to *Γ*. In case *α* < 1, the motion is considered subdiffusive (MacKintosh, 2012), which in this case could be caused by association of OCR-2 to stationary structures and *I* or by the restriction of OCR-2 motion due to other, slower moving proteins (Saxton, 1994). Application of this MSD analysis to the three example trajectories confirms the different modes of motility underlying these trajectories: the MSD of the apparently directed trajectory (*) scales roughly quadratically with time, that of the diffusive trajectory (+) linearly, and that of the hardly moving molecule (#) with a slope less than linear, indicative of subdiffusion.

**Figure 4:**
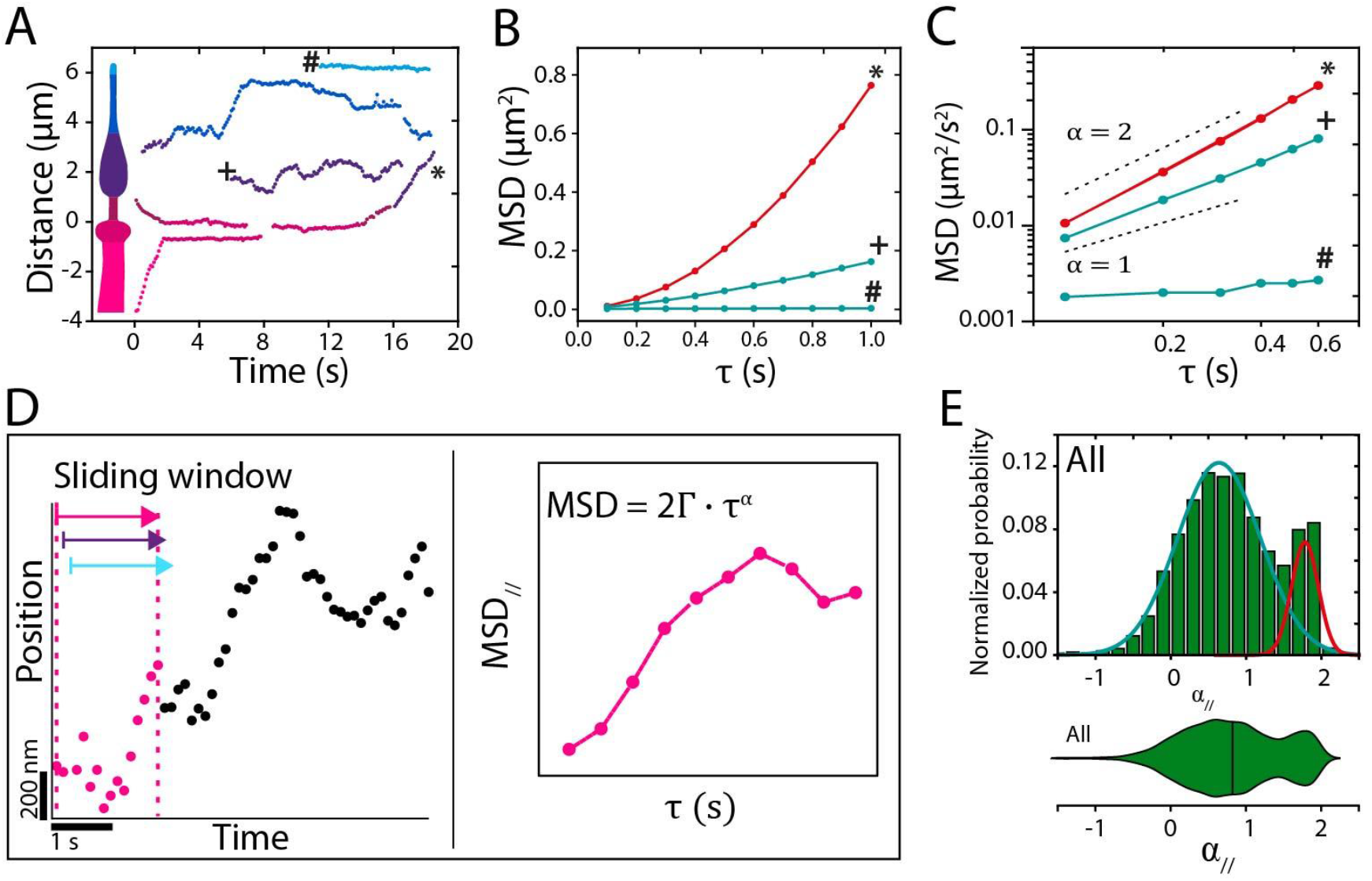
Analysis of single-molecule OCR-2 trajectories. **(A)** Example single-particle trajectories from the kymographs of figure 3A. **(B).** Mean squared displacement (MSD) of annotated trajectories in **(A),** demonstrating active transport (*), normal diffusion (+) and subdiffusion (#). (C) Log-log plot of the first six time points. (D) Cartoon explaining the single-molecule data analysis with sliding window. (E) Histogram and corresponding violin plot of normalized probability of *α*_//_ for all 6992 data points obtained from 23 nematodes. Colored lines represent two underlying Gaussian distributions.

The analysis used above was applied to user-selected (fragments of) trajectories with different length. In order to obtain unbiased insight from all 185 trajectories (in 23 worms) constituting 6992 individual localizations, we calculated the instantaneous MSD, using a sliding window of 15 consecutive locations, in accordance with previous work (Godin et al., 2017), yielding instantaneous values of Γ and *α*, both parallel and perpendicular to the cilium (Figure 4D and methods) (Danné, 2018). As a first characterization of the data obtained in this way, we plotted all *α*_//_ values in a histogram and corresponding violin plot (Fig 4E). Histogram and violin plot clearly show a bimodal distribution of *α*_//_ confirming that OCR-2 moves by a combination of directed and diffusive motion. A fit with two Gaussians to the histograms yields a diffusive fraction (84%) with average *α*_//_ of 0.66 (Full Width Half Maximum (FWHM): 0.56), and a directed motion fraction (16%) with average *α*_//_ of 1.78 (FWHM: 0.17). This indicates that the majority of motion of OCR-2 in the cilia is diffusive. Furthermore, in particular the diffusive fraction is broadly distributed, indicating heterogeneous diffusive behavior. In the remainder of this study we will use *α* = 1.4, the intersection between the two Gaussians fits, as the threshold to discriminate between diffusive and directed motion. This analysis indicates that instantaneous values of *α* can be readily obtained from single-molecule trajectories of OCR-2 in cilia and can provide quantitative insight in OCR-2 dynamics.

### A location-specific interplay of transport dynamics underlies the ciliary distribution of OCR-2

Next, we used the same dataset to obtain more quantitative insight into the motion of OCR-2 in the different ciliary segments. To obtain an overview of the location-specific motility properties of OCR-2, all *α*_//,⊥_ and *D*_//_ values, derived from the instantaneous MSD, were color coded for *a* and plotted in function of their location (Fig 5A). Even though the data used for these heat maps is derived from 23 different animals, the resulting shape resembles the shape of a single cilium, validating the robustness of our imaging and analysis. This representation furthermore highlights that OCR-2 motility parameters depend on the location in the cilium. To further quantify this, we represented the distributions of the *α*_//_ values in the different ciliary segments in violin plots (Figure 5B). The violin plots show that the distributions vary substantially between the segments. In the TZ, *α*_//_ values peak close to two; in the PS *α*_//_ values are widely distributed, peaking around 1; in the DS, two peaks can be observed, the major, broad one peaking around 0.5 and the minor one peaking slightly below 2; in the Tip, a single distribution around 0.5 can be seen. The results of Gaussian fits to these distributions (Figure S5) is represented in Figure 5C, which shows the fractional contributions for directed motion and diffusion in the different ciliary compartments and the corresponding average *α*_//_ values. Taken together, Figure 5B and C confirm, in a quantitative way, the initial indications obtained from the kymographs. In the TZ, OCR-2 moves mostly by directed transport; in the PS, OCR-2 moves mostly by normal diffusion in the membrane and only to a small extend via directed transport; in the DS, the fraction of OCR-2 moving by directed transport is slightly higher, while the rest moves subdiffusively; at the tip all motion is subdiffusive.

**Figure 5:**
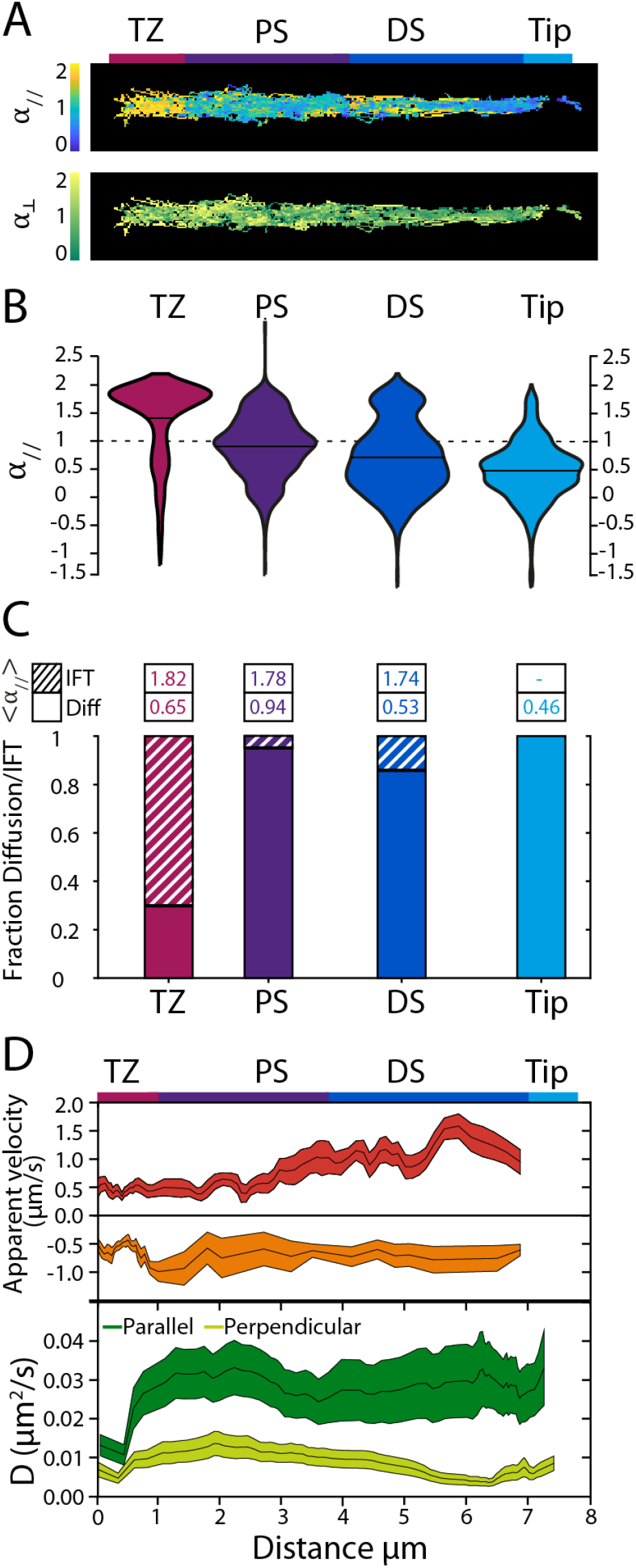
Analysis of single-molecule OCR-2 trajectories reveals location-specific motility. **(A)** Heat maps showing all data points color coded for *α*_//_ and *α*_⊥_. Scale bar: 1 μm. **(B)** Violin plots showing *α*_//_ for the TZ, PS, DS and Tip regions of the *C. elegans* phasmid cilia. (C) Relative fractions of molecules undergoing IFT or diffusive motion, with corresponding average *α*_//_ in the different ciliary segments. **(D)** Apparent velocities and diffusion coefficients of OCR-2 molecules in the phasmid cilia. Top: apparent anterograde and retrograde velocities determined for time points for which *α*_//_ ≥ 1.4, averaged over 10 displacements. Line thickness represents s.e.m.. Bottom: parallel and perpendicular diffusion coefficients of OCR-2 determined for time points for which 0.9 ≤ *α*_//,⊥_ ≤ 1.1, averaged locally over 60 displacements. Line thickness represents s.e.m..

We further analyzed our data by determining the apparent local velocities from (stretches of the) trajectories with *α*_//_ ≥ 1.4 using *v_app_* = (*x*(*t* + *dt*) – *x*(*t*))/*dt*, with *dt* the frame integration time (Holcman et al., 2018), averaging locally over 10 displacements (Figure 5D). Local apparent anterograde velocity compares very well to velocity profiles obtained before using bulk (Prevo et al., 2015) and singlemolecule fluorescence imaging (Oswald et al., 2018) of kinesin motors and other IFT components: with a velocity of about 0.5 μm/s close to the TZ, gradually increasing in PS, reaching the maximal value of more than 1 μm/s. This similarity of the anterograde OCR-2 velocity profiles presented here, compared to those obtained from the IFT components reported before, highlights the validity of our approach. Apparent retrograde velocities, however, agree less well with velocities reported before and are lower, in particular in the DS. We do not have a definitive explanation for this, but we note that the number of events showing directed minus end motion is substantially lower than for anterograde, potentially indicating that bouts of retrograde transport of OCR-2 are only short lived (compared to our sliding window of 15 frames, corresponding to 1.5 s). In agreement with this, the OCR-2 kymographs show only few straight retrograde events in contrast to anterograde events, which are far more evident.

We next looked into the values of the apparent diffusion coefficients parallel and perpendicular to the ciliary axis (*D*_//,⊥_), calculated from (segments of) trajectories with *α*_//.⊥_ ≤ 1.1, using *D*_//,⊥_ = (*r*_//,⊥_(*t* + *dt*) – *r*_//,⊥_(*t*))^2^/*dt* (with *r*_//,⊥_ = *x,y*) (Holcman et al., 2018), averaging locally over 60 displacements, Figure 5D. We made sure that *D*_⊥_ was determined for OCR-2 only when it was not actively transported by IFT by adding as constraint that only time windows were included with *α*_//_ < 1.4, in order to allow for a proper comparison of the diffusion coefficients in both directions. Using this approach, we found that *D*_//_ was rather constant, −0.03 μm^2^/s, along the cilium, except for the TZ, where it was −0.01 μm^2^/s. *D*_⊥_ was almost three times smaller than *D*_//_ −0.01 μm^2^/s, decreasing somewhat in the distal segment. In part, the lower *D*_⊥_ compared to *D*_//_ could be a consequence of our imaging approach, which effectively results in a 2-dimensional projection of a 3-dimensional object, making us blind for motion in the direction perpendicular to the image plane (which is one of the two axes perpendicular to the cilium). This could result in an underestimation of the diffusion coefficient in the direction perpendicular to the cilium, at most by only a factor of about 2 (Renner et al., 2011; Van Den Wildenberg et al., 2011). We speculate that the larger difference between the apparent *D*_//_ and *D*_⊥_ in DS and Tip might be caused by confinement, which could result in subdiffusive motion.

Taken together, single-molecule imaging reveals the extent of the location-specific diversity in motility modes of OCR-2 in *C. elegans* chemosensory cilia, ranging from active transport, normal diffusion, to subdiffusion. This combination of motility modes is crucial for establishing the steady-state distribution of OCR-2 over the cilium, which is essential for effective OCR-2 function, allowing the binding of aversive chemicals to receptors to provoke an avoidance reaction of the whole animal.

## Discussion

Exposed to repellent chemicals, *C. elegans* performs avoidance behavior. In this study, we have aimed to connect this whole-organism behavior to the dynamics of the molecular machinery inside the chemosensory cilia that sense the chemicals and trigger the subsequent processes that drive behavior. To this end, we have exposed *C. elegans* to aversive chemicals and imaged the distribution and dynamics of ciliary components using fluorescence microscopy. Our image sequences show that exposure to SDS results in a rapid (−10-20 s), reversible redistribution of tubulin, IFT-dynein, and OCR-2 out of the DS, about a minute after exposure to SDS. Furthermore, with single-molecule imaging, we have shown that the transmembrane calcium channel OCR-2 displays complex location-specific motility, ranging from subdiffusion to active transport by IFT. Together, these diverse motility modes help to establish OCR-2 distribution along the cilium and allow for its reversible redistribution.

Whereas the SDS-evoked avoidance response occurs almost instantaneously after SDS addition, the redistribution of ciliary components starts much later, about a minute after addition, suggesting that the redistribution is not an intrinsic part of the avoidance response. Laser-ablation experiments of the *C. elegans* phasmid dendrite and live imaging of IFT and calcium in *Chlamydomonas* have shown that the distribution of IFT components and IFT dynamics can be altered almost instantaneously (Collingridge et al., 2013; Mijalkovic et al., 2020). The cause of the delay in our SDS-addition experiments is hence not the inability of the cilia or IFT to react rapidly. Since in our experiments SDS exposure was prolonged, it is well possible that the delayed redistribution is part of a sensory habituation mechanism. Sensory habituation has been studied before in the ASH amphid chemosensory neurons in *C. elegans* (Hilliard et al., 2005). These experiments revealed that the calcium concentration in the ASH somas (and as a consequence the neuronal response) decreased on a −minute time scale during prolonged repellent exposure (Hilliard et al., 2005). The calcium response to repellents was also shown to decrease after short, consecutive stimuli. Taken together, this suggests that the reversible redistribution of ciliary components we observed a −minute after SDS exposure is indeed part of a sensory habituation mechanism, distinct from the avoidance response. Although questions remain about the exact function of the redistribution of ciliary components in chemosensing, it is well possible that the plasticity of the axoneme and ciliary composition could be involved in a mechanism increasing the dynamic range of chemosensing in order to minimize (prolonged) exposure to potentially dangerous compounds.

Distribution and motility of TPs involved in sensing have been studied in the cilia of other model systems, in particular in primary cilia of cultured mammalian cells (Huang et al., 2007; Milenkovic et al., 2015; Qin et al., 2005; Weiss et al., 2019; Ye et al., 2013). The advantage of using a multicellular organism such as *C. elegans* as a model organism is that, in principle, the whole cascade from sensory membrane protein, to secondary messengers, neuronal action and finally behavior can be studied in the same model. The price one has to pay is that high-sensitivity imaging is hampered by signal loss due to scattering and optical aberrations caused by imaging relatively deep into the tissue of the animal. Using our approach, it works well for the phasmid cilia in the tail of the nematode, but not for the amphid cilia in the head, because of more out-of-focus background fluorescence, higher autofluorescence of the surrounding tissue and substantially more movement of the animal. Imaging primary cilia of mammalian cultured cells poses, in this regard, substantially less challenges.

It is important to note that, although *C. elegans* chemosensory cilia are highly homologous to mammalian primary cilia, important differences exist. Whereas the primary cilia of cultured mammalian cells are slightly shorter and consist of an axoneme with doublet microtubules along almost their entire length, *C. elegans* cilia consist of an axoneme with doublet microtubules only until about halfway, with microtubule singlets extending towards the ciliary tip in the DS. Furthermore, *C. elegans* chemosensory cilia are located inside the amphid and phasmid channels with only the tip sticking out into the environment, while mammalian cilia are completely exposed. This might explain why OCR-2 in *C. elegans* chemosensory cilia accumulates at the tip, whilst GPCRs like Somatostatin Receptor 3 (SSTR3) and Smoothened, in mammalian primary cilia are more evenly distributed along the length of the cilia (Milenkovic et al., 2015; Ye et al., 2013).

Despite these fundamental differences in ciliary properties, our findings on the motility of OCR-2 in *C. elegans* phasmid cilia qualitatively agree with those of TPs in primary cilia of cultured mammalian cells. In *C. elegans we* found that OCR-2 motility is mostly due to diffusion and only less than 20% by active transport, IFT. Also in mammalian primary cilia, GPCRs have been found to be mostly diffusing, and only partly moved by IFT (−32-34% (Ye et al., 2013) and −1% (Milenkovic et al., 2015) IFT for Smoothened; −13-27% for SSTR3 (Ye et al., 2013); −4% for Patchedl (Weiss et al., 2019)). Remarkably, diffusion coefficients obtained for mammalian cilia are substantially larger (SSTR3: −0.25 μm^2^/s using FRAP (Ye et al., 2013); Smoothened −0.26 μm¾, using single-molecule tracking (Milenkovic et al., 2015); PTCHI: −0.1 μm^2^/s, PTCHI using single-molecule tracking (Weiss et al., 2019)) than what we observed in *C. elegans* phasmid cilia (−0.03 μm^2^/s). It is unclear whether these are actual differences or caused by the differences in measurement and analysis approaches, some being better in discriminating active transport from diffusion. On the other hand, a lower diffusion coefficient could be caused by a larger size of the transmembrane section of the calcium channel OCR-2 compared to the mammalian GPCRs, or by species-specific differences in ciliary membrane viscosity, potentially caused by protein crowding (Oswald et al., 2016). Notwithstanding these qualitative differences, our results and those in mammalian primary cilia indicate that the combination of diffusive and active transport motility modes of TPs, and their ability to connect and disconnect to underlying IFT trains, is a conserved mechanism for the motility of TPs in cilia. More than in these previous studies on mammalian primary cilia, our studies in *C. elegans* phasmid cilia, which are longer and have structurally different domains, have revealed that TP motility in cilia is segment dependent: in the TZ, OCR-2 is mostly transported directionally by IFT; in the PS, normal diffusion is prevalent; in the DS, subdiffusion is most common, with noticeably more directed transport than in the PS; in the tip, OCR-2 motility is rather low, mostly subdiffusive. These location-dependent motility modes and parameters most likely underlie the steady-state distribution of OCR-2 along the phasmid cilia.

What would be the molecular basis for this location-dependent motility? Most likely there are two distinct processes underlying this behavior. The first process is a location-dependent connection of OCR-2 with the IFT machinery. The interaction of OCR-2 and other TPs to IFT trains has been shown to be mediated by the BBSome protein complex, which is essential for chemotaxis (Blacque et al., 2004). Recent cryo-EM studies of the BBSome have revealed molecular details of this interaction, a negatively charged cleft on the BBSome surface that is likely involved in the binding of cargo proteins (Klink et al., 2019). In another study it has been suggested that the GTPase ARL6/BBS-3 is activated when it interacts specifically with the BBSome, allowing for the connection between IFT-trains and transmembrane cargoes (Chou et al., 2019). This is in agreement with our results, which suggest that this interaction is tightly regulated, being persistent in the TZ, and far more intermittent in the rest of the cilium. This transient connection between IFT machinery and OCR-2 might explain the differences in ensemble and SM kymographs we have observed. Ensemble kymographs hinted at OCR-2 undergoing IFT from base to tip, while singlemolecule kymographs showed far less regular, more saltatory motion. The ensemble IFT trajectories most likely are the result of multiple OCR-2 independently attaching to and detaching from a particular IFT train. Further studies will be required to better understand the interaction of TPs and the IFT machinery and how this is locally regulated.

The second process underlying the location-dependent motility is the location-dependent (sub)diffusion. We have shown that OCR-2, when not connected to the IFT machinery, diffuses normally (with *a* ≈ 1) in the PS, while it moves subdiffusively in the DS and Tip (with *a ≪* 1). Most likely, subdiffusion in the parallel direction is caused by an increased density of OCR-2 or other TPs (crowding), and/or static structural proteins, all hampering the motion of OCR-2 in the membrane. In addition, differences in membrane composition and curvature could also play a role, especially in the perpendicular direction in regions where the average distance explored during subdiffusion is close to the local diameter of the cilium. To further elucidate the molecular mechanisms inside the cilia underlying avoidance behavior, experiments will be required with a higher level of control over behavioral cues. In particular microfluidic devices could be employed to accurately control the exact instant and duration of a chemical stimulus (Chronis et al., 2007). This approach might allow imaging IFT at the single-molecule level, which is not feasible with our current approach. In addition, functional imaging approaches could be employed, including calcium probes to sense neuronal activity or probes that sense other secondary messengers including cAMP and cGMP (Imaging et al., 2019; Jiang et al., 2019).

In conclusion, we have shown that sensitive fluorescence microscopy is an invaluable tool to study how whole-organism behavior is connected to single-protein dynamics. Investigating the molecular machinery underlying development and function of cilia provides important insights into mechanisms involved in key life processes such as sensing, signal transduction, limb patterning, and human disorders collectively named ciliopathies (Iii et al., 2018; Tobin and Beales, 2009).

## Materials and methods

### *C. elegans* strains

Strains used in this study were generated using MOSCI insertions and CRISPR/Cas9 genome editing and are listed in Table S1 (Frøkjær-Jensen et al., 2008; Paix et al., 2017). Primers and guide RNA’s used to make the CRISPR/Cas9 and extrachromosomal array strains are listed in Table S2. Maintenance of *C. elegans* was performed using standard procedures (Brenner, 1974).

### Fluorescence microscopy

Fluorescence microscopy was performed as described previously (van Krugten and Peterman, 2018). In short, *C. elegans* young adult hermaphrodites were immobilized using 5 mM levamisole and mounted between a microscope slide with a 2% (W/V) agarose pad, and a coverslip. The worms were subsequently imaged on our custom-build wide-fìeld epifluorescence microscope with 100 ms exposure time, and gently photo bleached to obtain single-molecule sensitivity.

### Repellent addition experiments

Young adult hermaphrodite *C. elegans* were sedated as above, but mounted on a microscope slide with a central opening (AMIL Technologies). Imaging was performed as above. During the acquisition, 5 μl 0.1% (W/V) sodium dodecyl sulfate (SDS) in Ml3 was pipetted through the central opening, which was then cover with a 22 x 22 mm cover glass to prevent desiccation. The concentration of repellent at the worms was likely at least one order of magnitude lower than that what was added (Davies et al., 2003).

### Data analysis

Kymographs were generated from the image sequences using the open source ImageJ plugin KymographClear and stand-alone program KymographDirect (Mangeol et al., 2016). Single-molecule tracking was performed as described previously, using custom-written MATLAB (MathWorks) routines using a linking algorithm (Jaqaman et al., 2008; Prevo et al., 2015). To be able to define and distinguish movement parallel and perpendicular to the ciliary axoneme, trajectory coordinates were transposed to a hand-drawn spline using the custom-written MATLAB script. In order to distinguish anterograde and retrograde velocities of OCR-2, we calculated the sign of the difference between two consecutives parallel positions *sign*(*x*(*t* + *dt*) – *x*(*t*)). If the sign is positive, the directed motion is anterograde.

### Repellent experiments

#### Drift correction

Drift coming from the movement of the worms was first corrected using the Fiji plugin TrackMate with the LoG (Laplacian of Gaussian) detector option (Tinevez et al., 2017). This was used to find the location of a centroid of a immobile bright spot in the worm, which could be used as reference point. Using the position of this spot in each time point of the raw movie (M1), we generated a new movie (M2) corrected by the drift. However, this method is not robust enough due to the precision of localization of the Gaussian fit. In order to minimize the residual drift, we generated kymographs of movie M2 using the Fiji plugin KymographClear and extracted the position of the base using the intensity contrast in the kymograph (Mangeol et al., 2016). This second correction of the drift was then used to calculate the intensity in the different parts of the cilia.

#### Correction for spontaneous photobleaching

To correct the intensity values in the different ciliary segments for spontaneous photobleaching, the intensity values before repellent addition (*t* < *t*_0_) were fitted by a decreasing exponential curve (*A exp*(–*t/τ*) + *C*). The values of the intensity at each time of the movie were then multiplied by the exponential part *exp*((*t* + *t*_0_)/*τ*).

#### Averaging over multiple cilia

For TBB-4::EGFP and XBX-l::EGFP, the intensity values were corrected for spontaneous photobleaching and normalized by the values in the different parts at the beginning of the movies. These normalized intensities were then averaged (n = 8 for TBB-4 and n = 7 for XBX-1). For OCR-2, the intensities in the different ciliary segments were normalized by the sum of the intensity of the whole cilia. These normalized intensities were then averaged (n = 8).

## Supporting information

Movie S1: Redistribution of IFT-dynein upon repellent exposure.

Movie S2: Single-molecule imaging of SRB-6.

Movie S3: Single-molecule imaging of OCR-2.

## Acknowledgements

We thank J. Mijalkovic for helpful discussion and support, and Laurent Cognet and Antoine Godin (University of Bordeaux) for help and discussions regarding the analysis of single-molecule trajectories. We acknowledge financial support from the Netherlands Organisation for Scientific Research (NWO) via a Foundation for Fundamental Research on Matter (FOM) program grant (“The Signal is the Noise”) and from the European Research Council under the European Union’s Horizon 2020 research and innovation programme (Grant agreement no. 788363; ?ITSCIL”).

## Author Contributions

Conceptualization and Methodology, J.v.K. and E.J.G.P.; Investigation, JvK; Formal Analysis, J.v.K. and N.D.; Resources, J.v.K. and E.J.G.P.; Writing - Original Draft, J.v.K. and E.J.G.P.; Writing - Review & Editing, J.v.K., N.D. and E.J.G.P.; Funding Acquisition and Supervision, E.J.G.P.

## Supplemental information

### Nematodes experience prolonged exposure to SDS during repellent experiments

To investigate whether the time course of SDS concentration around the imaged worms might be the cause of the delay, we added fluorescein in an identical way as SDS and measured its intensity over time. Following an immediate increase of fluorescein intensity, which explains the immediate avoidance response, a plateau was reached after about 2 minutes. This indicates that a low concentration is sufficient to evoke an avoidance response.

**Figure S1:**
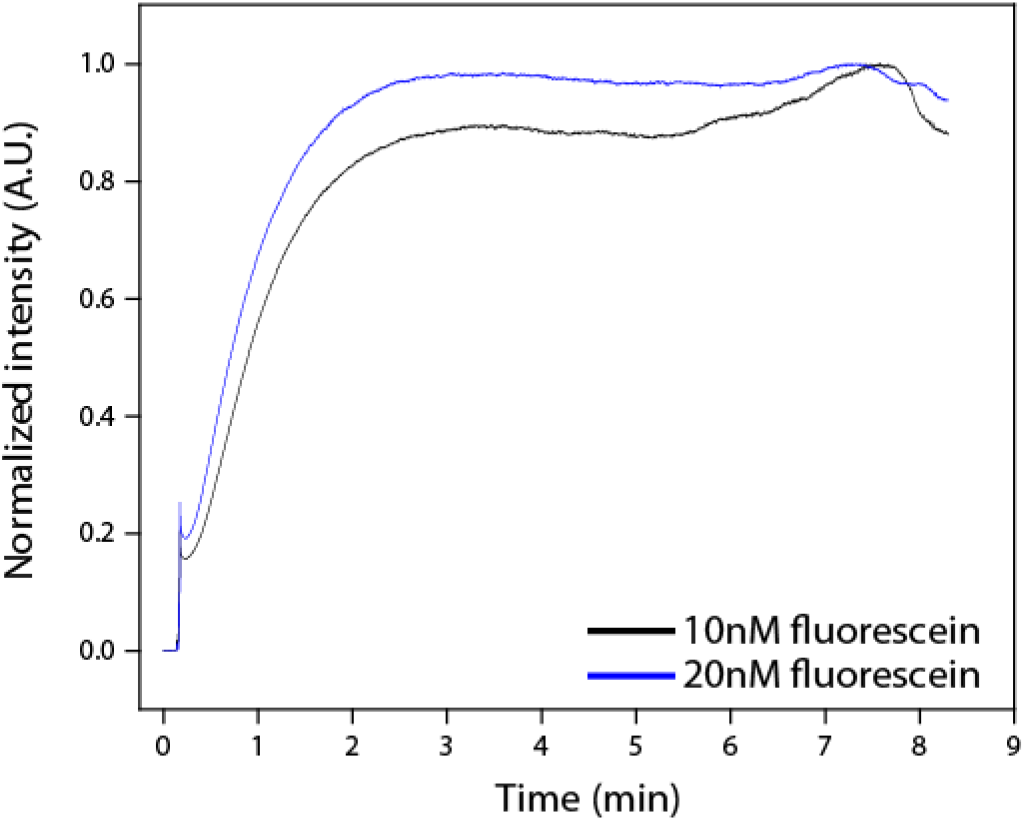
Intensity of fluorescein over time, after addition to sample. A 5 μl droplet of fluorescein (of 10nM and 20 nM) was added to an agarose pad on a cover slip, and the intensity measured over time.

**Figure S2:**
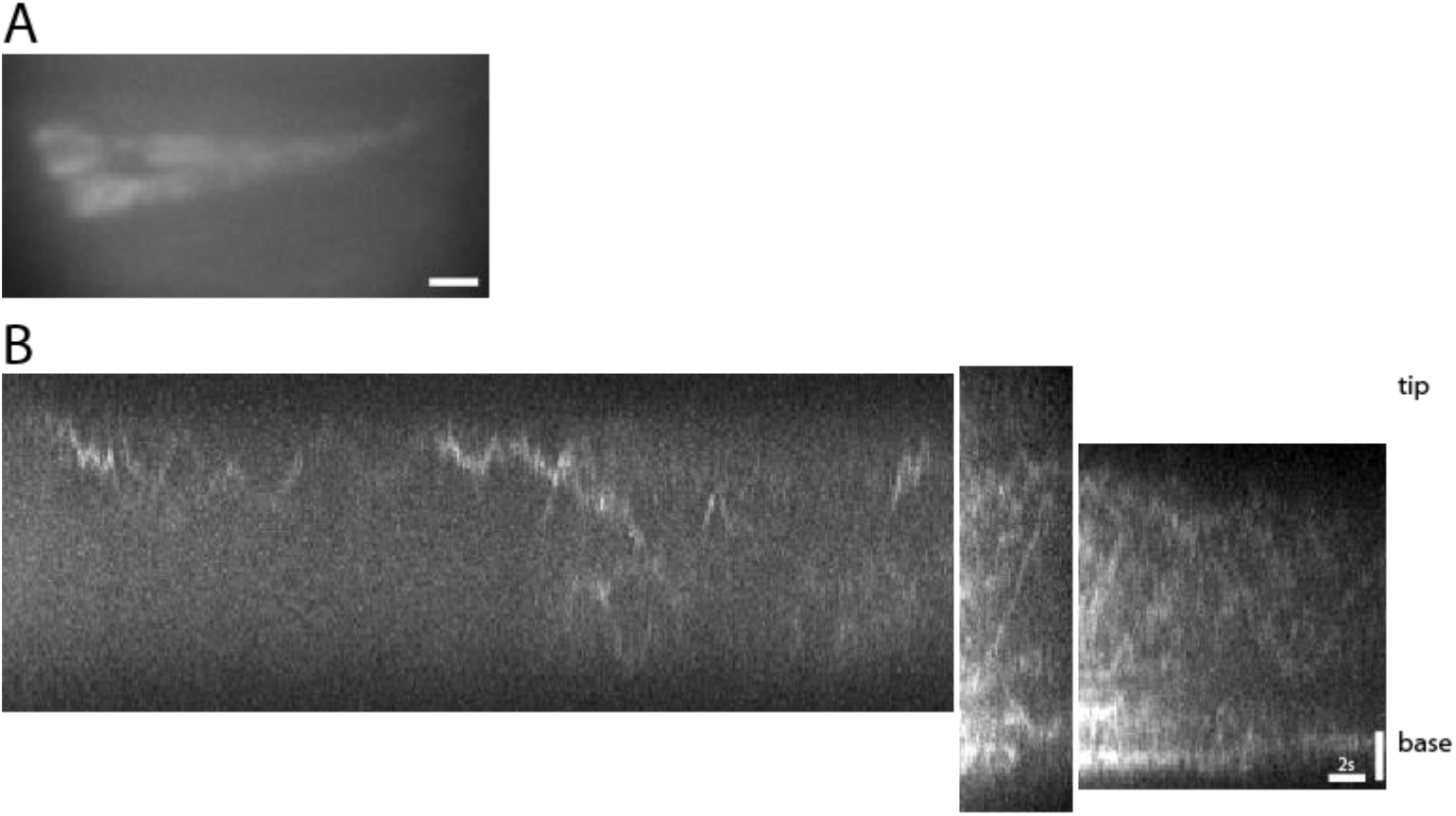
Low expression levels of GPCR SRB-6 in phasmid cilia. (A) Time-averaged image sequence shows the ciliary distribution of SRB-6::EGFP is comparable to that of OCR-2::EGFP (Scale bar 1 μm). (B) Kymographs showing salutatory movement and active transport of SRB-6 in the phasmid cilia (horizontal: time, 2 s; vertical: distance, scale bar 2 μm). Related to Movie S2.

**Figure S3:**
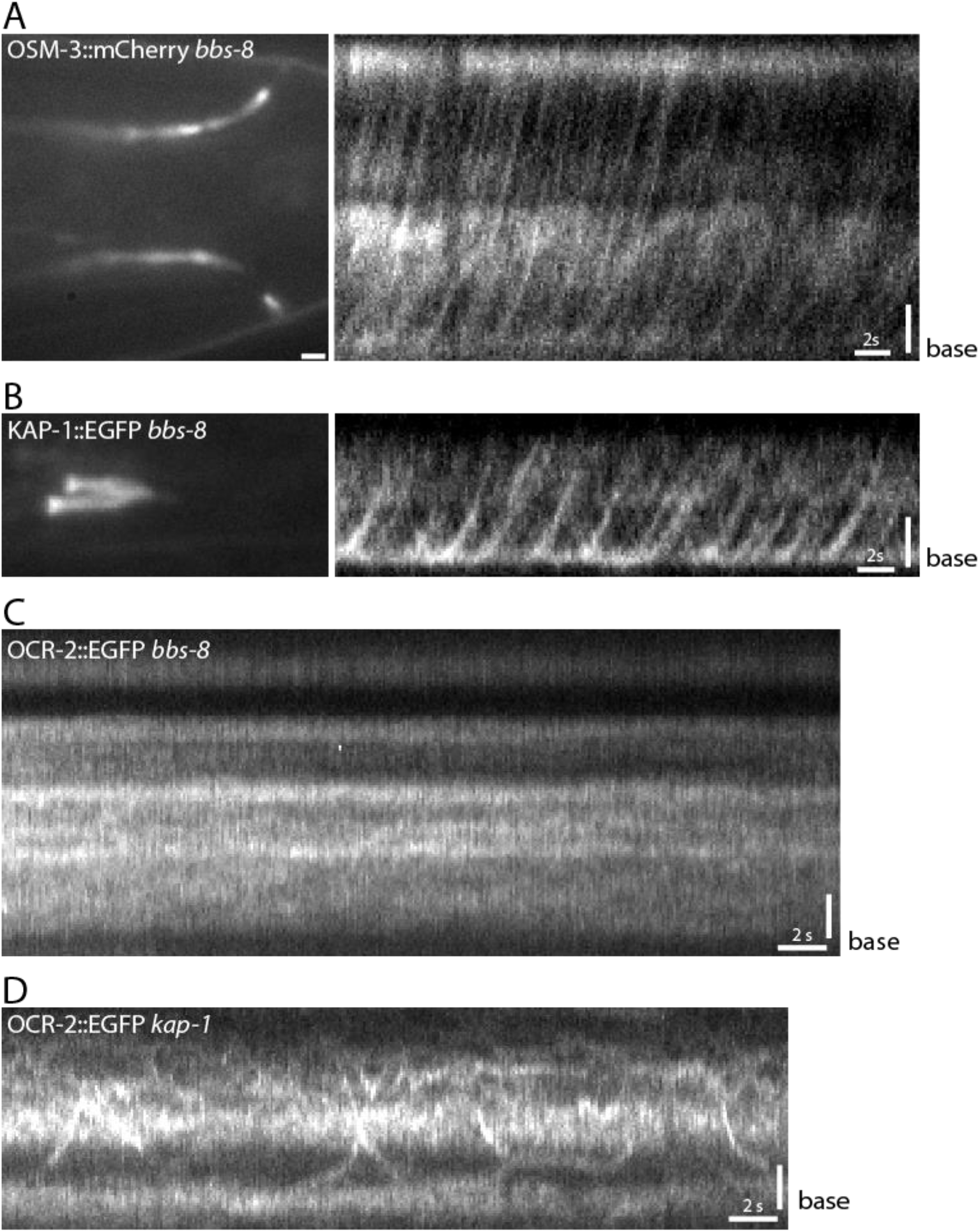
Functional IFT in cilia lacking functional BBSome. Time-average image sequences and kymographs of OSM-3::EGFP (A) and KAP-l::EGFP (B) in *bbs-8* background show functional IFT, despite being chemotaxis deficient. These data indicate that although the distribution of OCR-2 in *bbs-8* worms is drastically different, the cilia itself are still intact, and IFT is functional, though not coupled to OCR-2. (C) Kymograph of OCR-2::EGFP in *bbs-8* background, showing no IFT movements of OCR-2. (D) Kymograph of OCR-2::EGFP in kap-1 mutant background (horizontal: time, 2 s; vertical: scale bar 1 μm).

### OCR-2 is transported towards the cilia via the trans-golgi network

During the experiments to collect data for the time-averaged fluorescence image sequences, bright OCR-2 foci moving in the dendrite towards the PCMC were observed. As protein synthesis of ciliary components occurs in the soma of the sensory neuron, roughly 100 μm away from the cilium, we hypothesized these foci were vesicles of OCR-2 transported via the trans-golgi network. To confirm this, we generated a strain over expressing the vesicle marker RAB-8::mScarlet and endogenously labeled OCR-2 (Kaplan et al., 2010). Indeed, dual-color fluorescence imaging showed overlapping foci for OCR-2::EGFP and RAB-8::mScarlet. Furthermore, the velocity of the vesicles, measured by kymograph analysis, was comparable to that of cytoplasmic dynein.

**Figure S4:**
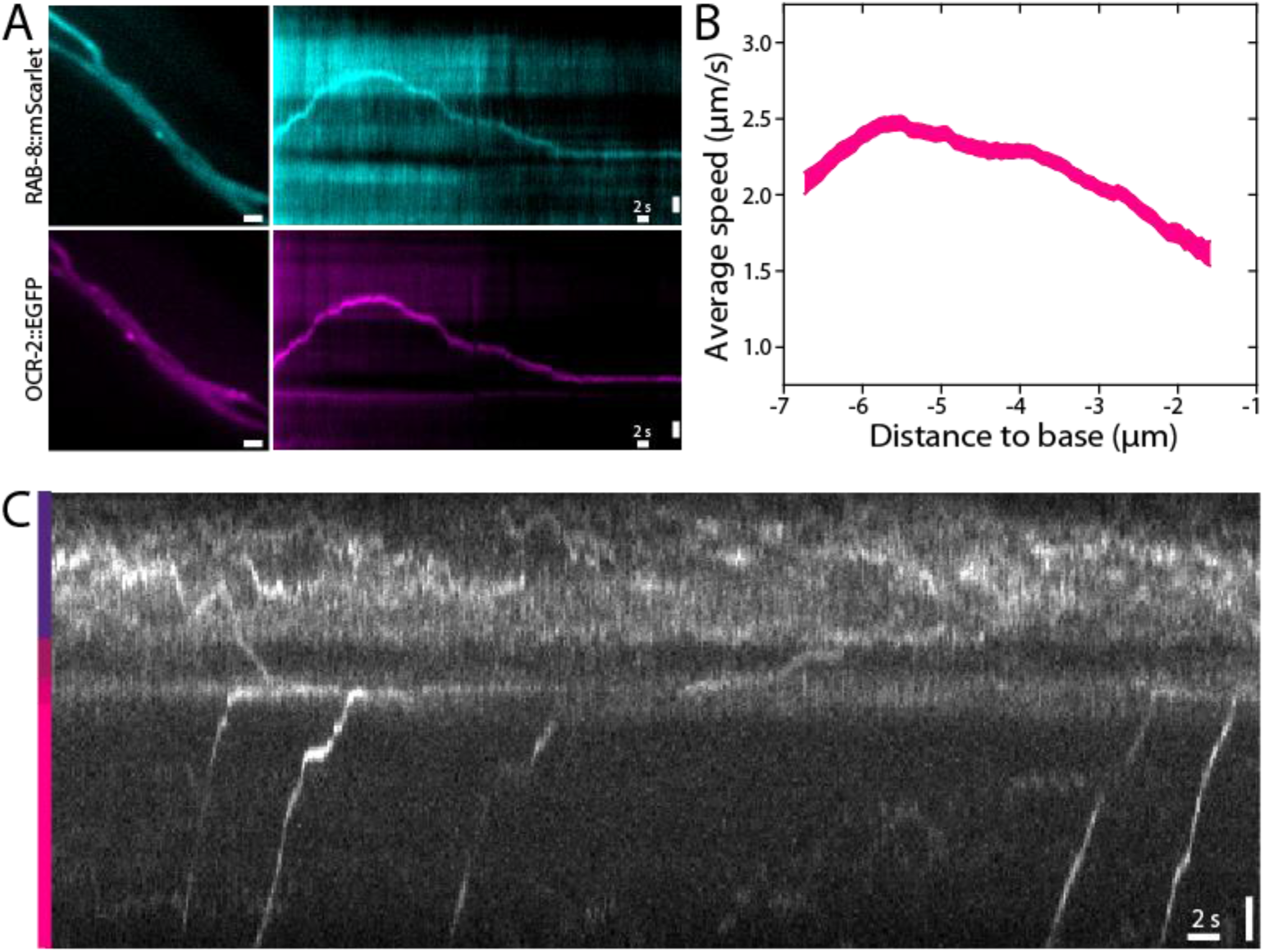
Dendritic transport of OCR-2 towards phasmid cilia. (A) Co-localization, as shown in a frame of the movie (left) and on a kymograph of the dendrite (right), of vesicle marker RAB-8 (cyan, top) and OCR-2 (magenta, bottom) in foci travelling in the phasmid neuron dendrite. (B) Average speed of OCR-2::EGFP foci traveling towards the cilium. (C) Kymograph showing OCR-2::EGFP foci arriving in the PCMC. Here, the signal gradually decreases, indicating docking and diffusion of OCR-2 with and in the membrane of the PCMC. A retrograde and anterograde TZ crossing can also be observed. Color bar on the left indicates the dendrite (bottom), PCMC, TZ and proximal segment (top) according to the diagram of figure 1.

**Figure S5:**
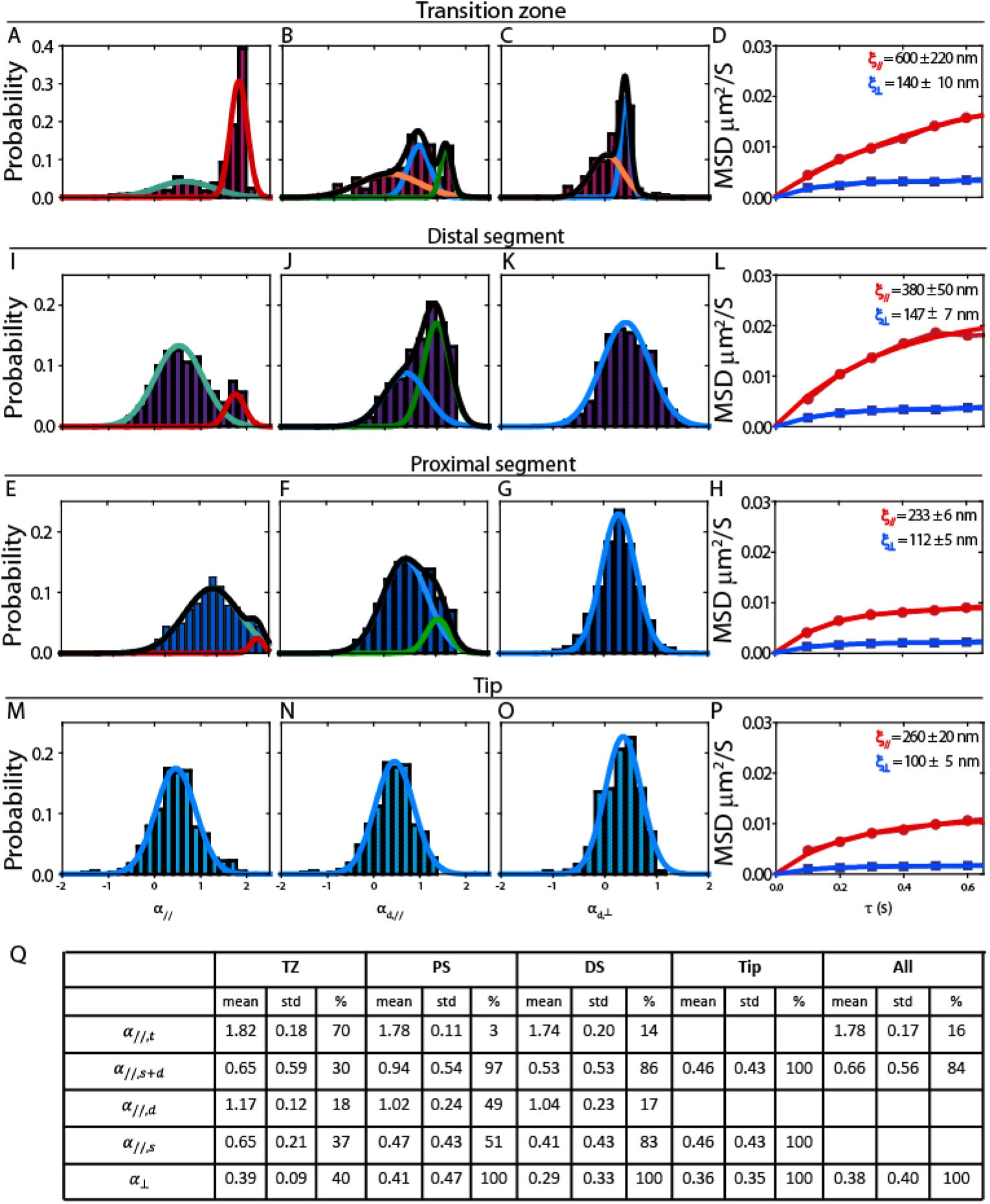
Summary of *α* values and extraction of the sizes explored during subdiffusion. (A, E, I and M) Histograms of *α*_//_ values in TZ, PS, DS and Tip respectively from all the trajectories. Straight curves indicate Gaussian fits of histogram showing transported and subdiffusion fractions, (B, F, J, N) Histograms of *α*_//_ values when *α*_//_ < 1.4 in TZ, PS, DS and Tip respectively. Straight curves indicate Gaussian fits of histogram showing diffusion and subdiffusion fractions, (C, G, K, O) Histograms of *α*_⊥_ values when *α*_//_ < 1.4 in TZ, PS, DS and Tip respectively. Straight curves indicate Gaussian fits of histogram showing diffusion and subdiffusion fractions, (D, H, L, P) Mean Square Displacement in function of the lag time in the parallel and perpendicular directions plotted for lower than unity values of *α*_//_ (blue circles) and *α*_⊥_ (red squares) respectively (with a range ± 0.05). Blue and Red curve indicated fits of MSD with the equation 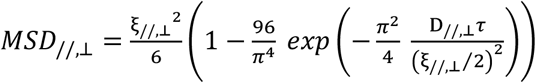. Values of size explored by OCR-2 (*ξ*_//,⊥_ are exrracred from fits. Table showing all *α*_//,⊥_ values extracted from Gaussian fits of histograms (A-C, E-G, I-K, M-O).

### Supplemental tables

**Supplemental table 1:**
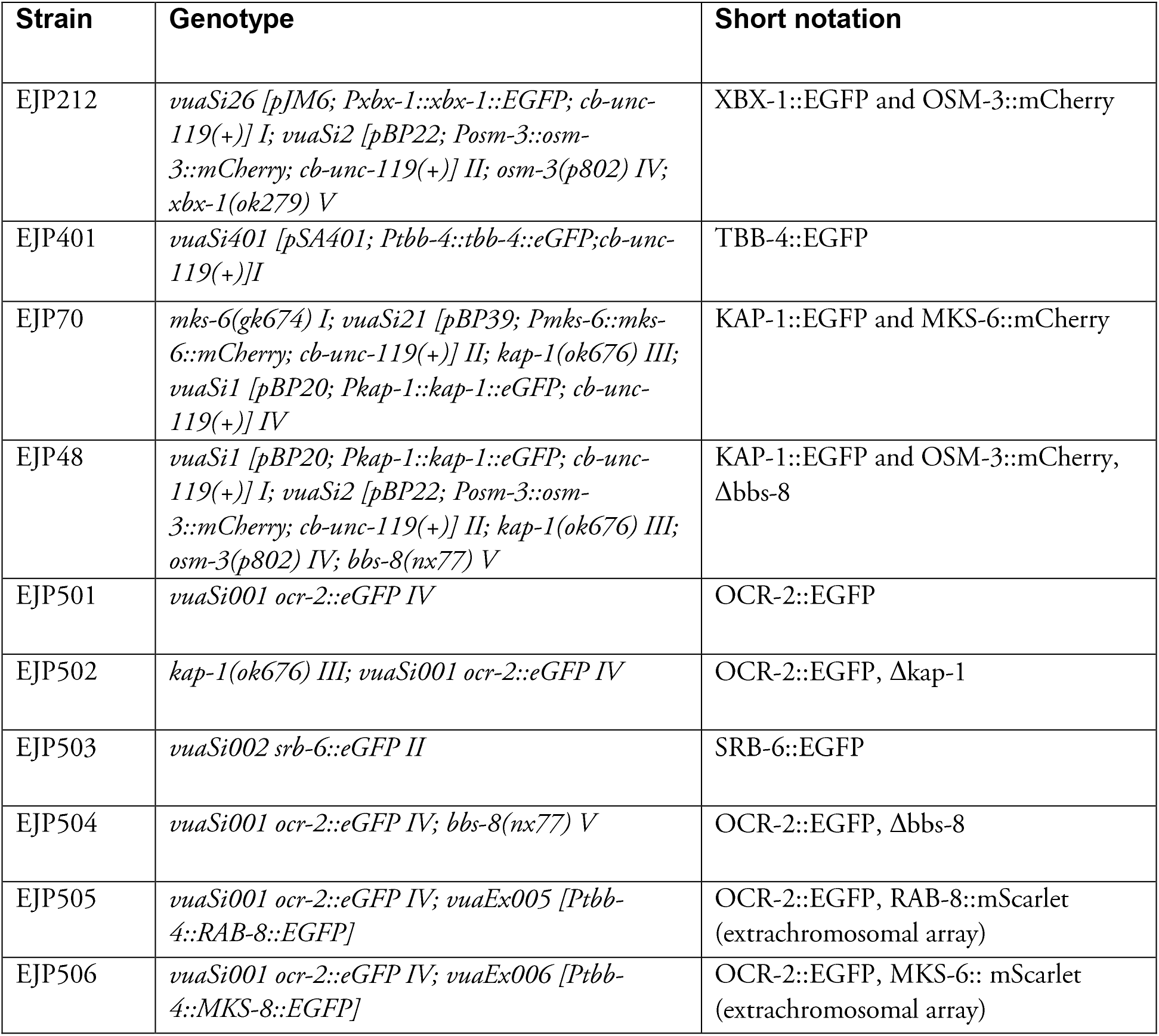
Strains used in this study.

**Supplemental table 2:**
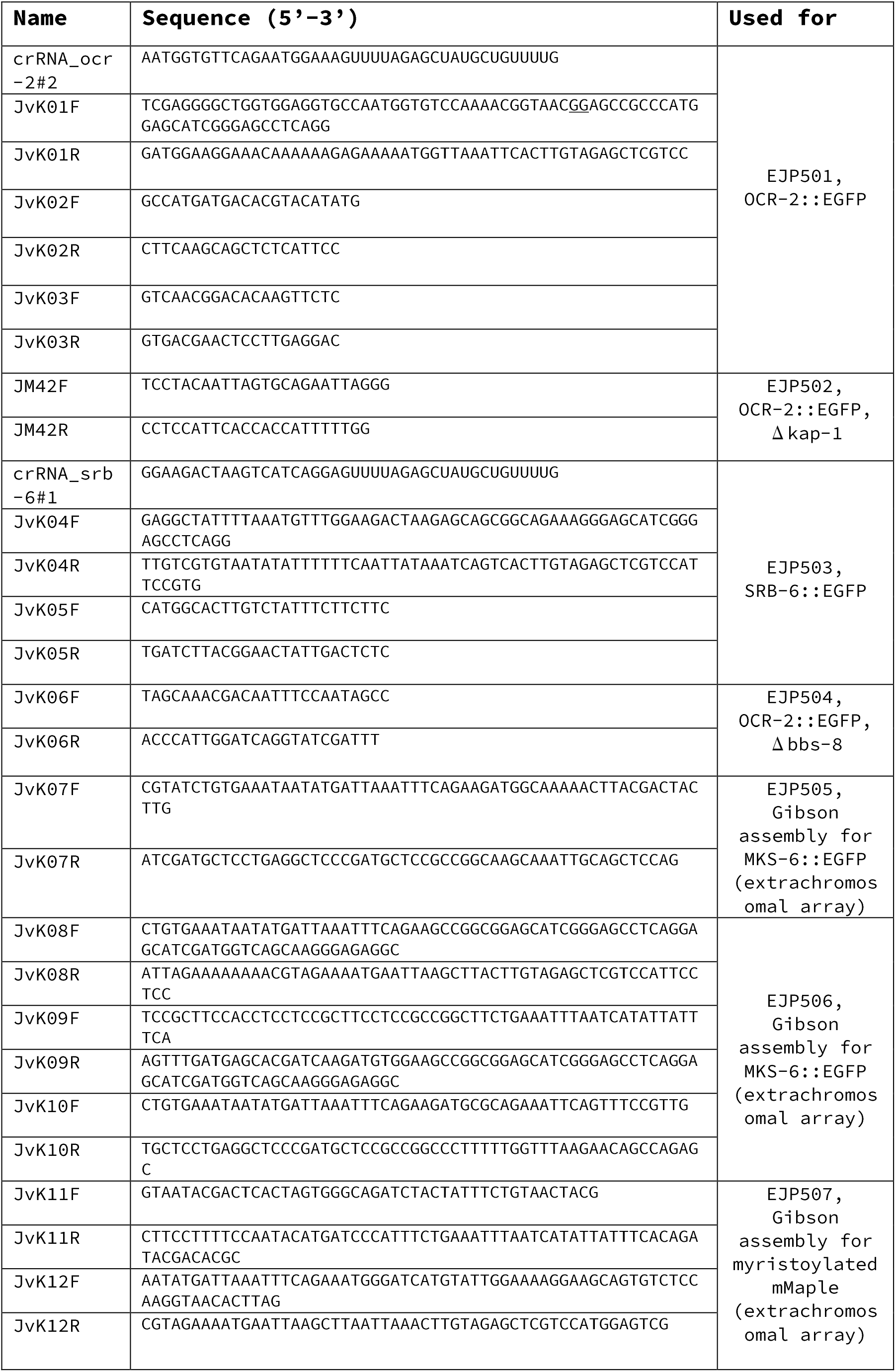
Primers used in this study.

### Supplementary movies

**Movie SI: Redistribution of IFT-dynein upon repellent exposure.** Fluorescence microscopy image sequence showing the reversible redistribution of IFT-dynein (XBX-l::EGFP) in the distal segments of the phasmid chemosensory cilia of *C. elegans* after the addition of a droplet of 5 μl 0.1% (W/V) SDS. Related to Figure 1.

**Movie S2: Single-molecule imaging of SRB-6.** Single-particle fluorescence image sequences of endogenously labeled SRB-6, showing its low expression level in the phasmid cilia and mostly diffusive and salutatory motility.

**Movie S3: Single-molecule imaging of OCR-2.** Collage of example single-molecule image sequences of OCR-2::EGFP, demonstrating the diversity in OCR-2 motility. Related to Figure 3A.

